# Gill developmental program in the teleost mandibular arch

**DOI:** 10.1101/2022.03.15.484461

**Authors:** Mathi Thiruppathy, J. Andrew Gillis, J. Gage Crump

## Abstract

Whereas no known living vertebrate possesses gills derived from the jaw-forming mandibular arch, it has been proposed that the jaw arose through modifications of an ancestral mandibular gill. Here we show that the zebrafish pseudobranch, which regulates blood pressure in the eye, develops from mandibular arch mesenchyme and first pouch epithelia and shares gene expression, enhancer utilization, and developmental *gata3* dependence with the gills. Combined with work in chondrichthyans, our findings in a teleost fish point to the presence of a mandibular pseudobranch with serial homology to gills in the last common ancestor of jawed vertebrates, consistent with a gill origin of vertebrate jaws.

## Introduction

Gills are the major sites of respiration in fishes. They are composed of a highly branched system of primary and secondary filaments, housing blood vessels, a distinct type of cellular filament cartilage, pillar cells (specialized endothelial cells), and epithelial cells maintaining ionic balance. In teleost gills, two rows of filaments are anchored to a prominent gill bar skeleton. Both the filaments and supportive gill bars develop from the embryonic pharyngeal arches that consist of mesenchyme of neural crest and mesoderm origin and epithelia of endodermal and ectodermal origin (Fabian et al., 2022; Mongera et al., 2013). The third through sixth arches generate gills in most fishes, and in cartilaginous fishes the second (hyoid) arch forms a hemibranch (one row of gill filaments). A classical theory for the origin of jaws posits that an ancestral gill support skeleton in the mandibular arch was repurposed for jaw function (Gegenbaur et al., 1878). Fossil evidence for ancestral vertebrates with a mandibular gill is, however, scant: exceptional soft tissue preservation of *Metaspriggina walcotti* from the Cambrian Burgess Shale had suggested a cartilaginous gill bar in the presumptive mandibular arch, yet gill filaments were not observed (Morris and Caron, 2014).

The pseudobranch is an epithelial structure located just behind the eye of most fishes that regulates ocular blood pressure. While it shares an anatomical resemblance to gill filaments, its embryonic arch origins remain debated (Miyashita, 2016). Our recent work in little skate points to a mandibular arch origin in chondrichthyans (Hirschberger and Gillis, 2021), which form one branch of jawed vertebrates. Whether the mandibular origin of the pseudobranch is conserved across vertebrates, including bony fishes, remained unknown, as well as whether the pseudobranch can be considered a serial homolog of the gills. Through lineage tracing and genetic analyses in zebrafish, we infer that the pseudobranch is likely a mandibular arch-derived structure in the last common ancestor of jawed vertebrates and represents a serial homolog of the gills.

## Results

In zebrafish, the pseudobranch is located anterior to the gill filaments and connected to the eye via the ophthalmic artery (Figure 1a,c) as described for other fishes (Laurent and Dunel-Erb, 1984). The pseudobranch appears in histological sections as a small bud behind the eye at 4 days post-fertilization (dpf) (Figure 1b, adapted from http://zfatlas.psu.edu/). Examination of *Sox10:Cre; acta2:Lox-BFP-Stop-Lox-dsRed* zebrafish shows this bud to be composed of a core of Cre-converted dsRed+ neural crest-derived cells ensheathed by unconverted BFP+ epithelia (Figure 1-figure supplement 1). The position of this bud corresponds to *kdrl:mCherry* labeling of a branch of the first aortic arch that likely gives rise to the ophthalmic artery (Figure 1-figure supplement 1). At 17 dpf, the pseudobranch is composed of five distinct filaments that resemble the primary gill filaments, with the five filaments merging to form a single pseudobranch by adult stages (90 dpf) (Figure 1b). Alcian Blue staining reveals that the adult pseudobranch contains five cartilage rods, reflecting the five fused filaments, with this cartilage resembling the specialized filament cartilage seen in the gills (Figure 1d) (Fabian et al., 2022).

**Figure 1.**
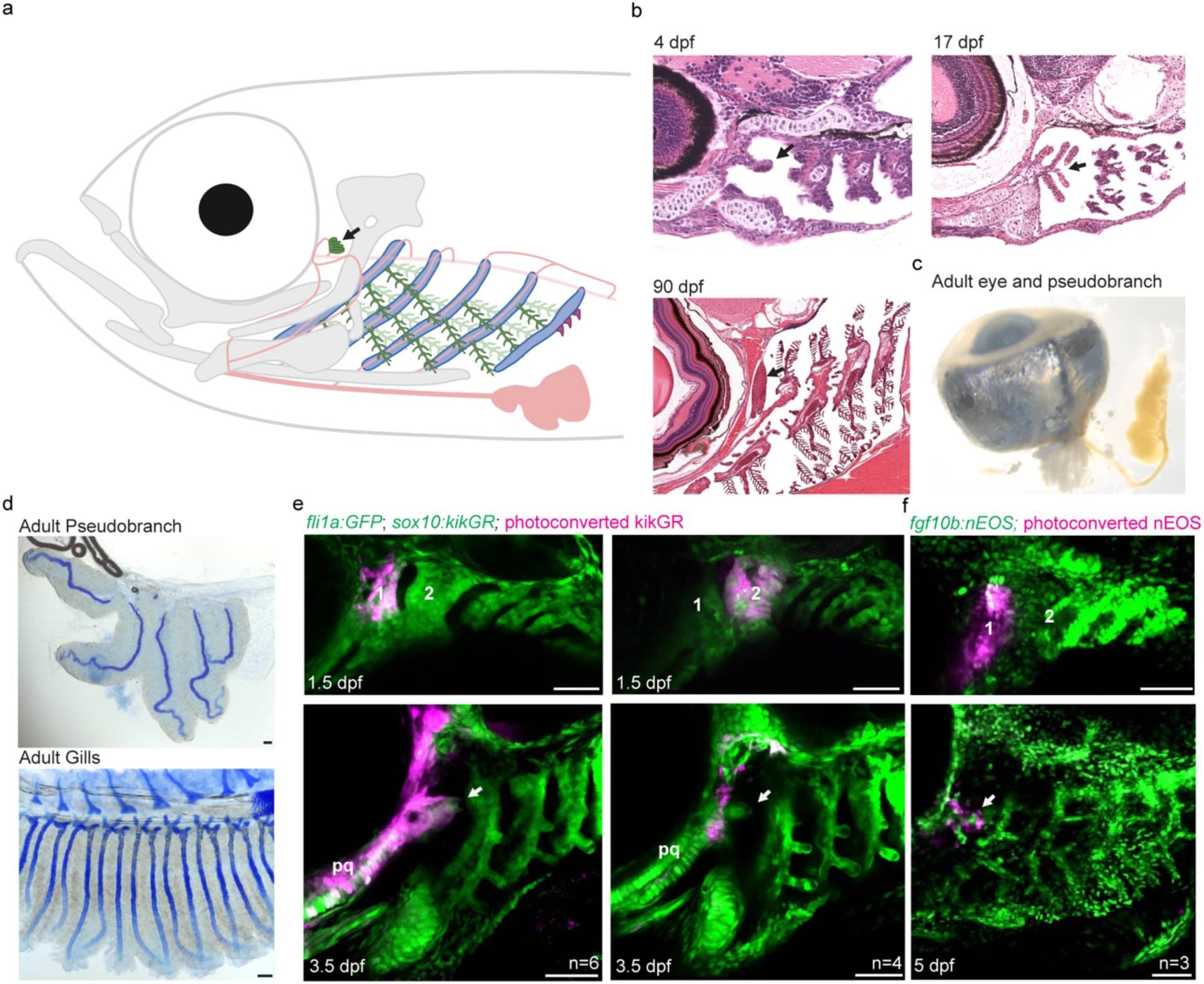
The zebrafish pseudobranch derives from mandibular arch mesenchyme and first pouch epithelia. **a**, Schematic showing the pseudobranch (arrows), gill filaments (branched green structures) connected to gill bars (blue), teeth (purple), vasculature (pink), and jaw support skeleton (grey). **b**, Hematoxylin and Eosin-stained sections show emergence of the pseudobranch bud at 5 dpf, five filaments at 17 dpf, and the fused pseudobranch at 90 dpf. **c**, Dissected adult pseudobranch shows the ophthalmic artery connecting it to the eye. **d**, Alcian staining shows five cartilage rods in the pseudobranch and similar cartilage in gill primary filaments. **e**, Photoconverted kikGR-expressing mesenchyme (red) from the dorsal first arch (numbered) at 1.5 dpf contributes to the palatoquadrate cartilage (pq) and pseudobranch mesenchyme (arrow) at 3.5 dpf. Photoconverted dorsal second arch cells do not contribute to the pseudobranch. In green, *fli1a*:GFP labels the vasculature and neural crest-derived mesenchyme for reference in green, and unconverted *sox10:kikGR* cells also labels mesenchyme. **e**. In *fgf10:nEOS* embryos, photoconversion of first pouch endoderm (numbered) at 1.5 dpf labels the pseudobranch epithelium (arrow) at 5 dpf. n numbers denote experimental replicates in which similar contributions were observed. Scale bars, 50 µm.

To determine from which arch the pseudobranch arises, we performed short-term lineage tracing using a photoconvertible *sox10:kikGR* reporter expressed in neural crest-derived mesenchyme. Photoconversion of dorsal first arch mesenchyme at 1.5 dpf labeled the pseudobranch mesenchymal bud at 3.5 dpf, as well as the palatoquadrate cartilage, a known first arch derivative; photoconversion of dorsal second arch mesenchyme did not label the pseudobranch (Figure 1e). To trace the epithelial origins of the pseudobranch, we performed short-term lineage tracing using *fgf10b:nEOS*, in which the photoconvertible nuclear-EOS protein is expressed in endodermal pouch epithelia (Figure 1-figure supplement 1). Photoconversion of first pouch endoderm at 1.5 dpf labeled pseudobranch epithelia at 5 dpf (Figure 1f), similar to labeling of first gill filament epithelia after photoconversion of third pouch endoderm (Figure 1-figure supplement 1). The pseudobranch therefore arises from mandibular arch mesenchyme and first pouch epithelia.

In skate, the pseudobranch and gills share expression of *foxl2, shh, gata3*, and *gcm2* (Hirschberger and Gillis, 2021). To test whether this reflects conserved gene regulatory mechanisms indicative of serial homology, we examined activity of several gill-specific enhancers (Fabian et al., 2022). At 5 dpf, the gata3-p1 enhancer drives GFP expression in the growing tips of both the pseudobranch and gill buds (Figure 2a). At 14 dpf, the ucmaa-p1 enhancer, active in gill filament but not hyaline cartilage in the face, drives GFP expression in both pseudobranch and gill filament cartilage (Figure 2b), as seen for endogenous expression of *ucmaa* (Figure 3-figure supplement 1). In our single-cell chromatin accessibility analysis of neural crest-derived cells (Fabian et al., 2022), we also identified an *irx5a* proximal enhancer selectively accessible in pillar cells, a specialized type of endothelial cell in the gill secondary filaments (Figure 3-figure supplement 2). At 13, 20, 60 dpf and one-year-old adult fish, the irx5a-p1 enhancer drives GFP expression in pillar cells of the pseudobranch and gills (Figure 2c; Figure 3-figure supplement 2). These findings show that the pseudobranch contains cells in common with both the primary and secondary filaments of gills, as well as conserved cis-regulatory architecture controlling gene expression.

**Figure 2.**
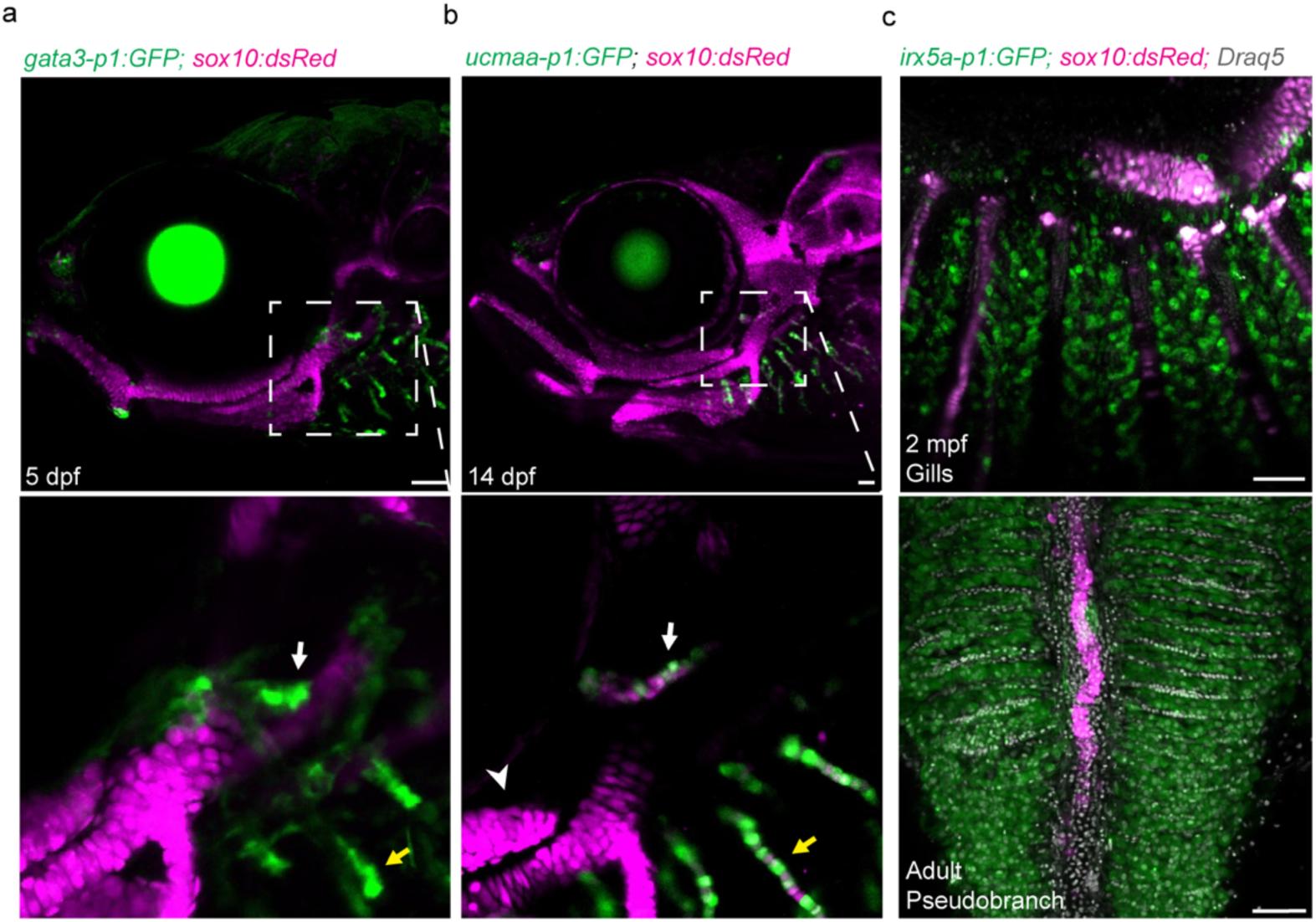
Conserved regulatory program for pseudobranch and gill development. **a-c**, In the pseudobranch (white arrows) and gill filaments (yellow arrows), *gata3-p1:GFP* labels growing buds, *ucmaa-p1:GFP* labels cellular cartilage (distinct from hyaline cartilage, arrowhead), and *irx5a-p1:GFP* labels pillar cells. *sox10:dsRed* labels cartilage for reference, and Draq5 labels nuclei in **c**. Magnified images are shown below for *gata3-p1:GFP* and *ucmaa-p1:GFP*. Scale bars, 50 µM.

Zebrafish mutant for *gata3* fails to form gill buds (Sheehan-Rooney et al., 2013), and single-cell chromatin accessibility analysis of neural crest-derived cells had implicated Gata3 and Gata2a in development of gill filament cell type differentiation (Fabian et al., 2022). We find that *gata3* and *gata2a* are prominently expressed in both the developing pseudobranch and gill buds at 3 and 5 dpf (Figure 3a; Figure 3-figure supplement 1). The pseudobranch is also much reduced in *gata3* mutants at 5 dpf, with fewer neural crest-derived cells labeled by *Sox10:Cre; acta2:Lox-BFP-Stop-Lox-dsRed* or *gata3-p1:GFP* contributing to both the pseudobranch and gills (Figure 3b,c). Similar genetic dependency of the pseudobranch and gills further supports serial homology.

**Figure 3.**
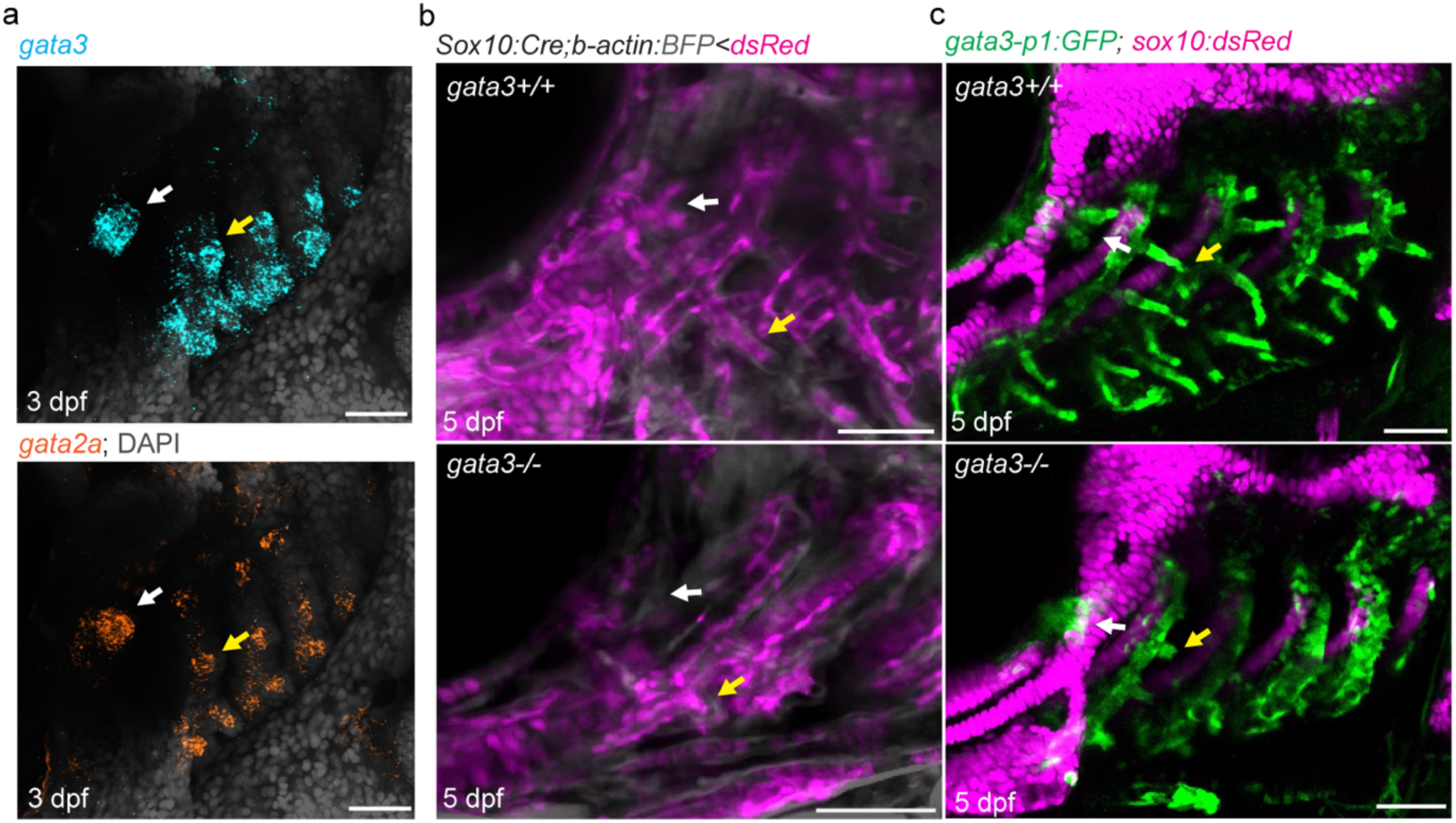
Pseudobranch and gill development require *gata3* function. **a**, Similar expression of *gata3* and *gata2a* in developing pseudobranch (arrow) and gill regions. **b**, *Sox10:Cre; acta2:Stop-BFP-Stop-dsRed* labels Cre-converted dsRed+ neural crest-derived mesenchyme (magenta) and unconverted BFP+ epithelia (grey). **c**, *gata3-p1:GFP* labels pseudobranch and gill filament buds, and *sox10:dsRed* labels cartilage. For both **b** and **c**, 3/3 *gata3* mutants displayed reduced formation of the pseudobranch (white arrows) and gill filaments (yellow arrows), compared to 3 controls each. Scale bars, 50 µM.

## Discussion

Our finding that the last common ancestor of jawed vertebrates likely had a mandibular gill-like pseudobranch provides further plausibility to the model that jaws evolved from a mandibular gill. Whereas our data clearly point to the filament systems of the pseudobranch and gills being homologous, the major skeletal bars supporting the jaws and gills (not to be confused with the gill filament cartilage) appear to develop largely independently from the filaments. Unlike the gill filaments, the zebrafish pseudobranch is not attached to a major skeletal bar. Conversely, the skeletal bars derived from the seventh arch of zebrafish lack gill filaments and instead anchor pharyngeal teeth, and the rostral-most gill bar reported for *Metaspriggina walcotti* lacked gill filaments (Morris and Caron, 2014). In addition, *gata3* loss affects the pseudobranch and gill filaments but not the gill bars (Sheehan-Rooney et al., 2013). It is therefore possible that, rather than the pseudobranch evolving from an ancestral mandibular gill whose gill bar was transformed into the jaw skeleton, the pseudobranch arose independently from the jaw by co-option of a gill filament developmental program. While we demonstrate gill-like developmental potential of the mandibular arch across vertebrates, whether an ancestral mandibular gill bar gave rise to vertebrate jaws awaits more definitive fossil evidence.

## Acknowledgements

We thank Megan Matsutani for fish care, Peter Fabian for dissection of the adult pseudobranch, and Keith Cheng, Jean Copper, and Daniel Vanselow for providing the original images related to Figure 1b.

## Methods

### Zebrafish lines

The Institutional Animal Care and Use Committee of the University of Southern California approved all animal experiments (Protocol 20771). Zebrafish lines include *Tg(fli1a:eGFP)*^*y1*^ (Lawson and Weinstein, 2002); *Tg(sox10:kikGR)*^*el2*^, *Tg(ucmaa_p1:GFP, cryaa:Cerulean)*^*el851*^, *Tg(gata3_p1:GFP, cryaa:Cerulean)*^*el858*^, and *Tg(fgf10b:nEOS)*^*el865*^ (Fabian et al., 2022); *Tg(−5.0sox17:Cre-ERT2,myl7:DsRed)*^*sid1Tg*^ (Hockman et al., 2017); *Tg(−3.5ubb:loxP-STOP-loxP-mCherry)* (Fabian et al., 2020); *Tg(Mmu.Sox10-Mmu.Fos:Cre)*^*zf384*^ (Kague et al., 2012); *Tg(actab2:loxP-BFP-STOP-loxP-dsRed)*^*sd27*^ (Kobayashi et al., 2014); and *gata3*^*b1075*^ (Sheehan-Rooney et al., 2013). To generate *Tg(irx5a-p1:GFP, cryaa:Cerulean)*^*el859*^, we synthesized the intergenic peak associated with *irx5a* (chr7:35838071-35838577) using iDT gBlocks, cloned it into a modified pDest2AB2 construct containing the E1b minimal promoter, GFP, polyA, and the lens-specific *cryaa:Cerulean* marker using in-Fusion cloning (Takara Bio). We injected plasmid and Tol2 transposase RNA (5-10 ng/µL each) into one-cell stage zebrafish embryos and screened for founders at adulthood based on lens CFP expression in progeny. Two independent germline founders were identified that showed activity in gill pillar cells.

### Histology

Adult fish were fixed in 4% paraformaldehyde for 1 hour at 25°C followed by dissection of the gills and further fixation in 4% paraformaldehyde for 1 hour at 25°C. For pseudobranch dissection, adults were fixed in 4% paraformaldehyde at 4°C for 3 days prior to dissection. Alcian Blue staining was performed on whole tissue as previously described (Paul et al., 2016). Samples were imaged using a Leica DM2500 microscope. Image levels were adjusted in Adobe Illustrator.

### Photoconversion-based lineage tracing

To photoconvert mesenchyme in *sox10:kikGR*; *fli1a:GFP* fish at 1.5 dpf, we used the ROI function in ZEN software on a Zeiss LSM800 confocal microscope to expose dorsal first or second arch mesenchyme to UV light for 20 s. Imaging confirmed successful and specific photoconversion of kikGR from green to red fluorescence in the intended region. At 3.5 dpf, confocal imaging was used to assess contribution of photoconverted cells to pseudobranch mesenchyme. We included *fli1a:GFP* to help in identification of the pseudobranch bud. For *fgf10b:nEOS* photoconversion, we used the ROI function to expose nEOS-high expressing cells in the first or third pharyngeal pouches to UV light for 60 s, with immediate confocal imaging confirming intended photoconversion of nEOS from green to red fluorescence. At 5 dpf, confocal imaging was used to assess contribution of photoconverted cells to pseudobranch and gill epithelia. To confirm that nEOS-high expressing cells in the *fgf10b:nEOS* line were of endodermal original, we crossed these onto the *sox17:CreERT2; ubb:Stop-mCherry* transgenic background and treated embryos with 4-Hydroxytamoxifen (Sigma) at 6.5 hpf to induce Cre recombination. We then imaged on the confocal microscope at 1.5 dpf to visualize co-localization of nEOS and mCherry. All results were independently confirmed in at least three animals.

### In situ hybridization

We performed in situ hybridization on whole embryos at 3 and 5 dpf and on paraffin sections from adult zebrafish heads using RNAscope probes synthesized by Advanced Cell Diagnostics in channel 1 (*ucmaa, gata2a*) and channel 2 (*gata3*). Samples were prepared by fixation in 4% paraformaldehyde overnight. Embryos were dehydrated in methanol and stored overnight before proceeding with the RNAScope Assay for Whole Zebrafish Embryos as described in the manufacturer’s protocols. Following fixation, the pseudobranch was dissected and mounted in 0.2% agarose in molds. Once solidified, agarose chips containing the pseudobranch were cut out of the mold, dehydrated, embedded in paraffin, and 5 µm sections were collected using a Shandon Finesse Me+ microtome (cat. no. 77500102). Paraformaldehyde-fixed paraffin-embedded sections were deparaffinized, and the RNAscope Fluorescent Multiplex V2 Assay was performed according to manufacturer’s protocols using an ACD HybEZ Hybridization oven. In situ patterns were confirmed in at least three independent animals, with exception of the *ucmaa* in situ that was performed on three separate sections of the same animal.

### Imaging

Images of whole-mount or section fluorescent in situ hybridizations and live transgenic fish were captured on a Zeiss LSM800 confocal microscope using ZEN software. For adult imaging of the *irx5a-p1:GFP* reporter, whole animals were euthanized and the pseudobranch and gills dissected out. The tissue was stained with Draq5 nuclear dye (Abcam) for 20 m to help identify pillar cells. Reported expression patterns for enhancer lines were confirmed in at least five animals.

---

**Figure 1-figure supplement 1.**
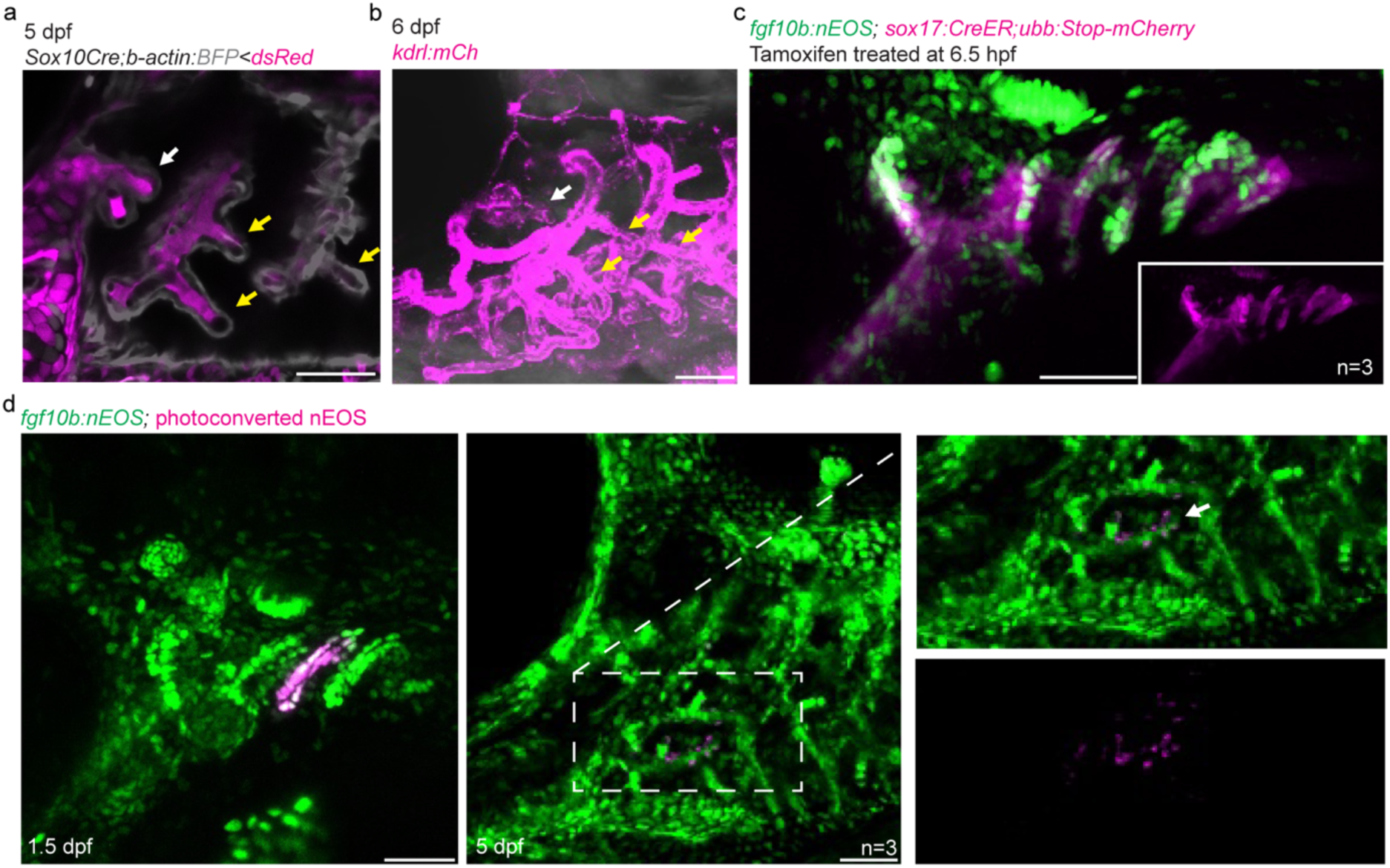
Development of zebrafish pseudobranch and lineage analysis of gill filament epithelia. **a**, In *Sox10:Cre; acta2:Stop-BFP-Stop-dsRed* fish at 5 dpf, the developing pseudobranch (white arrow) and gill buds (yellow arrows) consist of Cre-converted dsRed+ neural crest-derived mesenchyme (magenta) and unconverted BFP+ epithelia (grey). **b**, At 6 dpf, *kdrl:mCherry* labeling of vasculature reveals a branch of the first aortic arch in the position of the pseudobranch (white arrow), and branches of the posterior aortic arches in the positions of the gills (yellow arrows). **c**, Pharyngeal arch expression is seen in green for *fgf10b:nEOS*. Endoderm is labeled in red by adding 4OH-tamoxifen to *sox17:CreERT2; ubb:Stop-mCherry* fish at 6.5 hours post-fertilization to induce Cre recombination that removes the Stop cassette and allows mCherry expression (mCherry channel alone shown in inset). Co-localization shows *fgf10b:nEOS* expression in endodermal pouches. **d**, In *fgf10:nEOS* embryos, photoconversion of third pouch endoderm cells (numbered) at 1.5 dpf labels the first gill filament epithelium (arrow, boxed region magnified to right and shown in merged and red-only channels) at 5 dpf. Scale bars, 50 µM.

**Figure 3-figure supplement 1.**
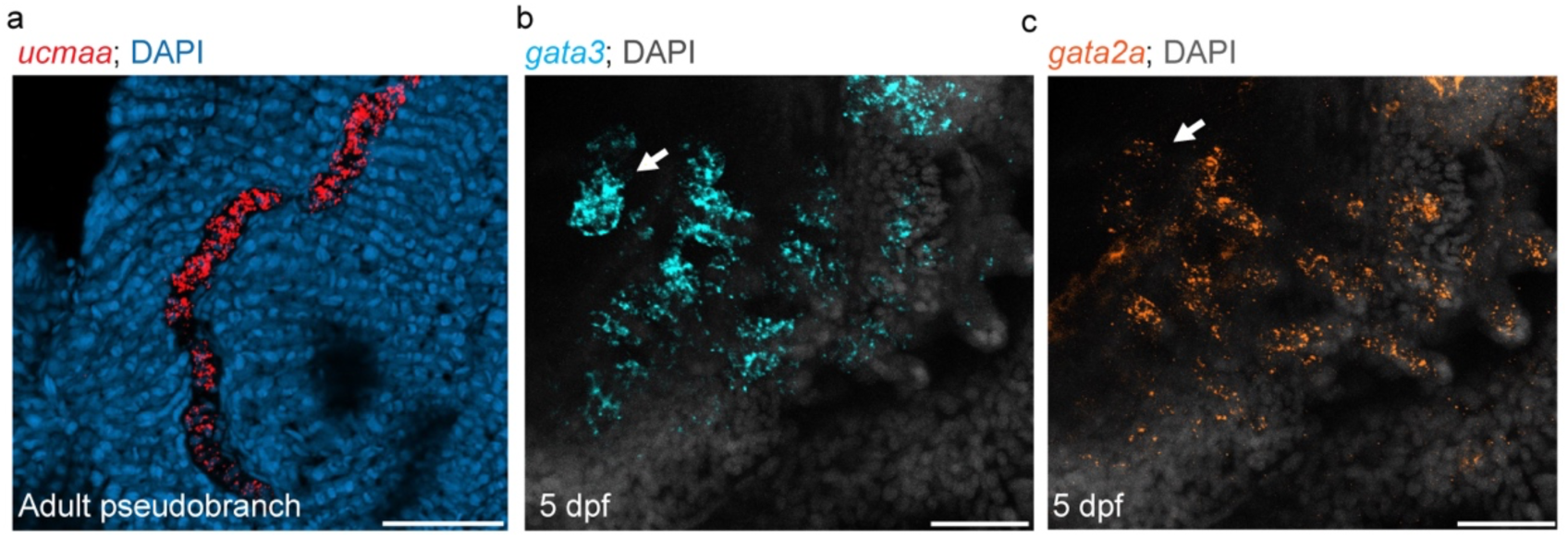
Pseudobranch shares gene expression with gill filaments. **a**, Expression of *ucmaa* in pseudobranch cartilage from one-year-old fish. **b**,**c**, Developing pseudobranch (arrows) and gill buds express *gata3* and *gata2a* at 5 dpf. DAPI labels nuclei in blue (**a**) or white (**b**,**c**). Scale bars, 50 µM.

**Figure 3-figure supplement 2.**
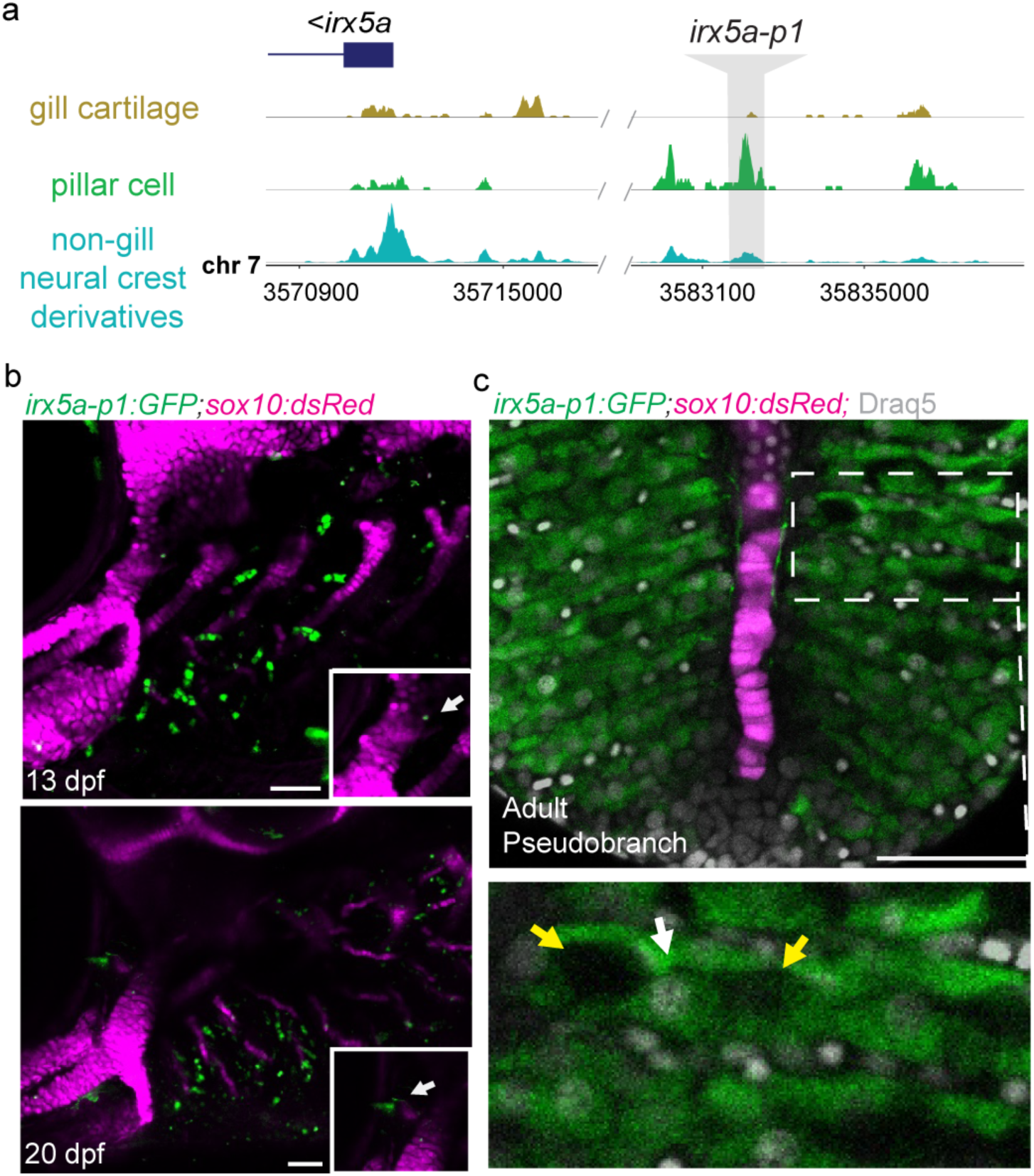
The pseudobranch shares *irx5a-p1* pillar cell enhancer activity with gill filaments. **a**, An intergenic irx5a-p1 region displays accessible chromatin specifically in pillar cells at 60 dpf. **b**, irx5a-p1 drives GFP expression in pillar cells of the developing gills and pseudobranch (inset, arrows). **c**,**d**, Dissected pseudobranch from a 60 dpf *irx5a-p1:GFP* adult shows GFP-positive pillar cells adjacent to the central core of filament cartilage. The boxed region magnified below depicts individual pillar cells (white arrow) interspersed by characteristic lacunae (yellow arrows). Cartilage is labeled by *sox10:dsRed* in **b** and **c**. Scale bars, 50 µM.

## References

Fabian, P., Tseng, K. C., Smeeton, J., Lancman, J. J., Dong, P. D. S., Cerny, R. and Crump, J. G. (2020). Lineage analysis reveals an endodermal contribution to the vertebrate pituitary. Science 370, 463–467.

Fabian, P., Tseng, K. C., Thiruppathy, M., Arata, C., Chen, H. J., Smeeton, J., Nelson, N. and Crump, J. G. (2022). Lifelong single-cell profiling of cranial neural crest diversification in zebrafish. Nat Commun 13, 13.

Gegenbaur, C., Bell, F. J. and Lankester, E. R. (1878). Elements of comparative anatomy. London,: Macmillan and Co.

Hirschberger, C. and Gillis, J. A. (2021). The pseudobranch of jawed vertebrates is a mandibular archderived gill. bioRxiv.

Hockman, D., Burns, A. J., Schlosser, G., Gates, K. P., Jevans, B., Mongera, A., Fisher, S., Unlu, G., Knapik, E. W. and Kaufman, C. K. (2017). Evolution of the hypoxia-sensitive cells involved in amniote respiratory reflexes. Elife 6, e21231.

Kague, E., Gallagher, M., Burke, S., Parsons, M., Franz-Odendaal, T. and Fisher, S. (2012). Skeletogenic fate of zebrafish cranial and trunk neural crest. PLoS One 7, e47394.

Kobayashi, I., Kobayashi-Sun, J., Kim, A. D., Pouget, C., Fujita, N., Suda, T. and Traver, D. (2014). Jam1a–Jam2a interactions regulate haematopoietic stem cell fate through Notch signalling. Nature 512, 319–323.

Laurent, P. and Dunel-Erb, S. (1984). The Pseudobranch: Morphology and Function. Fish Physiology 10, 285–323.

Lawson, N. D. and Weinstein, B. M. (2002). In vivo imaging of embryonic vascular development using transgenic zebrafish. Dev Biol 248, 307–318.

Miyashita, T. (2016). Fishing for jaws in early vertebrate evolution: a new hypothesis of mandibular confinement. Biol Rev Camb Philos Soc 91, 611–657.

Mongera, A., Singh, A. P., Levesque, M. P., Chen, Y. Y., Konstantinidis, P. and Nusslein-Volhard, C. (2013). Genetic lineage labeling in zebrafish uncovers novel neural crest contributions to the head, including gill pillar cells. Development 140, 916–925.

Morris, S. C. and Caron, J. B. (2014). A primitive fish from the Cambrian of North America. Nature 512, 419–422.

Paul, S., Schindler, S., Giovannone, D., de Millo Terrazzani, A., Mariani, F. V. and Crump, J. G. (2016). Ihha induces hybrid cartilage-bone cells during zebrafish jawbone regeneration. Development 143, 2066–2076.

Sheehan-Rooney, K., Swartz, M. E., Zhao, F., Liu, D. and Eberhart, J. K. (2013). Ahsa1 and Hsp90 activity confers more severe craniofacial phenotypes in a zebrafish model of hypoparathyroidism, sensorineural deafness and renal dysplasia (HDR). Dis Model Mech 6, 1285–1291.

